# Assembly-independent measurement of human Yq12 heterochromatin reveals population-scale variation and Yq12-derived RNA in testis

**DOI:** 10.64898/2026.11.11.765874

**Authors:** Mark Loftus, Pille Hallast, Gianni V. Martino, Gianluca Petris, Charles Lee, Miriam K. Konkel

## Abstract

Despite advances in long-read sequencing and telomere-to-telomere assembly, population-scale analysis of large satellite arrays remains challenging. Human Yq12 is a megabase-scale heterochromatic region composed primarily of *DYZ1*/HSat3A6 and *DYZ2*/HSat1B repeats, whose near-equal proportions are maintained despite extensive variation in total array length. Here, using complete Y-chromosome assemblies, we developed k-mer-based tools to estimate Yq12 size directly from short- and long-read sequencing data without requiring sample-specific complete assemblies. Across 1,597 males from the 1000 Genomes Project, Yq12 length ranged from ∼1 to 52 Mb (mean 26.5 Mb) and varied substantially within and between Y haplogroups. Phylogenetic analyses supported long-term constraint of total Yq12 length near an equilibrium of 26 Mb and strongly coupled evolution of *DYZ1* and *DYZ2* around near-parity composition. In 303 father-son pairs, fathers had longer estimated Yq12 regions in 70.6% of pairs, while sons more often showed *DYZ1*:*DYZ2* ratios closer to 1:1. Testis Iso-Seq identified transcripts containing *DYZ1* and *DYZ2* satellite sequences, while single-cell RNA-seq revealed their expression in haploid spermatids. These results establish a scalable framework for studying Yq12 population variation, intergenerational dynamics, and transcription, providing new opportunities to investigate how extreme satellite-array variation relates to Y-chromosome evolution and male germ-cell biology.

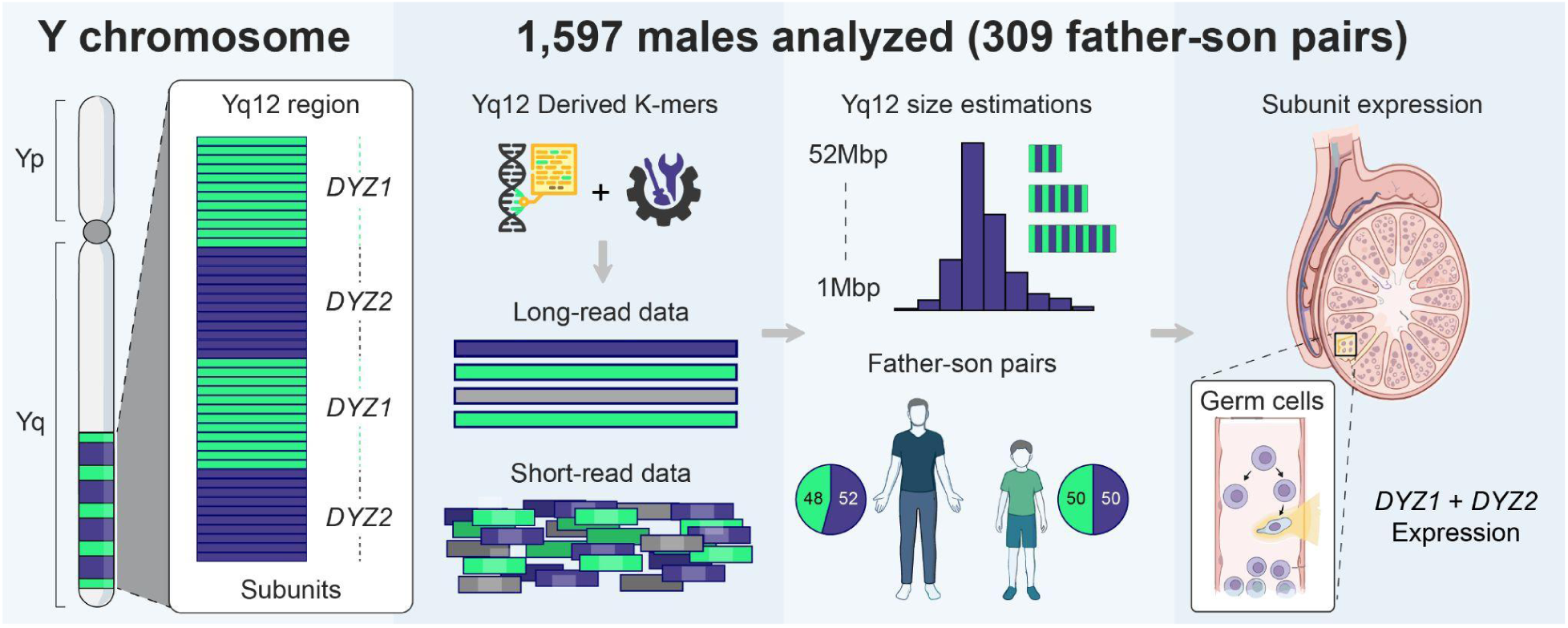

## Introduction

Satellite DNA forms megabase-scale domains at centromeres, pericentromeric regions, acrocentric short arms, and the distal long arm of the human Y chromosome. Its repetitive organization has historically prevented complete assembly and accurate population-scale measurement, leaving some of the largest components of the human genome underrepresented in reference sequences and genetic studies^1,2^. The Y chromosome is an extreme case: it was among the last human chromosomes to be completely assembled^3–5^, largely because most of its length comprises the non-recombining region of the Y (NRY), which contains extensive segmental duplications, ampliconic sequences, satellite arrays, and structural rearrangements^3,5,6^. Telomere-to-telomere and pangenome assemblies have since resolved much of the chromosome-specific organization and sequence composition of these regions while revealing substantial structural diversity among individuals^2,4–7^.

Among human chromosomes, the Y shows the greatest proportional variation in total length. Across 142 *de novo* assembled Y chromosomes, the total assembled chromosome length ranged from approximately 37.7 to 81.7 Mb, with a mean of 56.7 Mb^6^, although the full extent of human Y chromosome size variation is likely even greater. Much of this variability is the result of Yq12 size variation. The Yq12 region represents a human-specific megabase-scale block of tandemly repeated satellite DNA on the distal long arm of the chromosome (**Figure 1a**)^4,5^. Yq12 is composed primarily of two repeat families: *DYZ1*, also referred to as HSat3A6, which consists of approximately 3.5 kb repeat units that are enriched for simple and pentameric satellite sequences, and *DYZ2*, also known as HSat1B, which is composed of approximately 2.4 kb units comprising an AT-rich simple repeat fused to a truncated *Alu* element followed by HSat1 sequence^1,4,5,8^. These satellite sequences were first described in the 1970s^9,10^, with subsequent studies refining their genomic locations and sequence composition^11–14^. Together, these repeat units form long, alternating arrays that extend for tens of megabases (**Figure 1a**). Cytogenetic analyses have indicated that Yq12 can encompass between 29% and 54% of the metaphase Y chromosome length, with a median of 44%^15^, but these approaches provided only broad size estimates.

**Figure 1:**
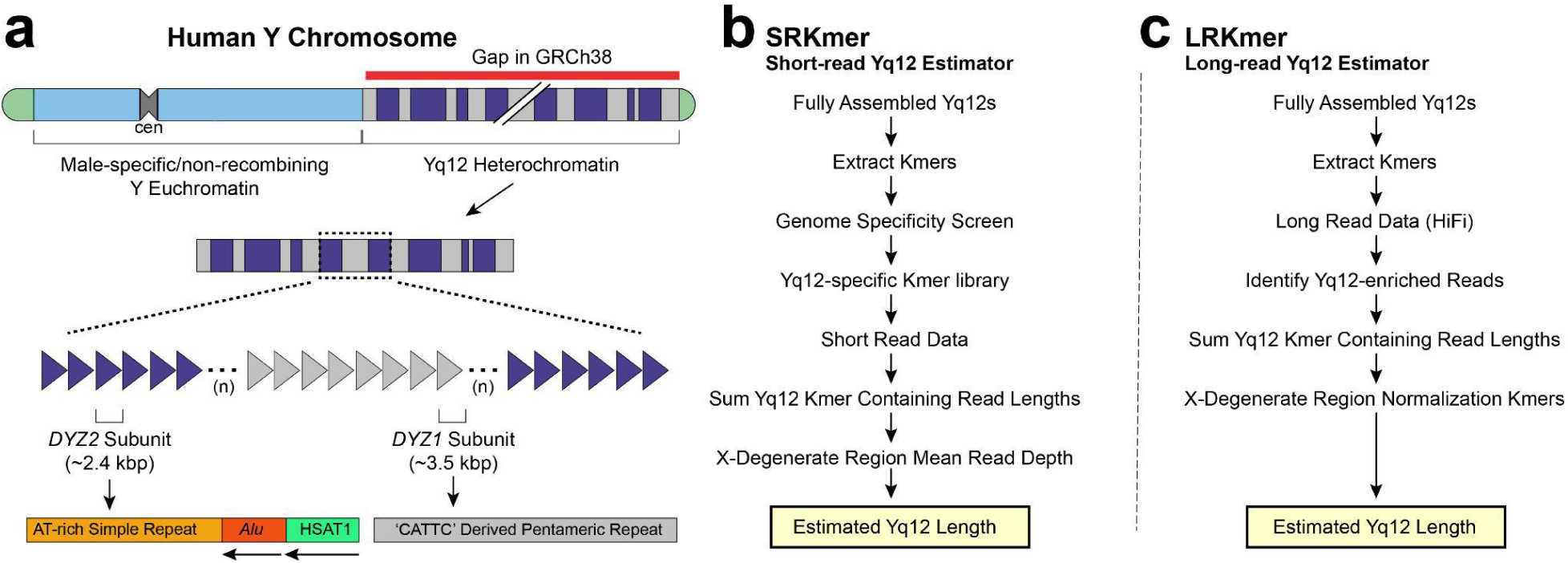
Structure of the human Yq12 heterochromatic region and overview of the Yq12 length-estimation pipelines. **a**) Schematic representation of the human Y chromosome highlighting the Yq12 heterochromatic region, which corresponds to a major gap in the GRCh38 reference assembly. Yq12 is composed predominantly of large tandem arrays of the *DYZ1* and *DYZ2* satellite repeats. b,c) Overview of the short-read k-mer (SRKmer) (**b**) and long-read k-mer (LRKmer) (**c**) Yq12 estimators. Both methods use k-mers derived from fully assembled Yq12 sequences to identify Yq12-derived reads and estimate Yq12 length from their cumulative sequence length. SRKmer uses a genome-wide-filtered library of Yq12-specific k-mers and normalizes by mean sequencing depth across Y-chromosome X-degenerate regions, whereas LRKmer identifies Yq12-enriched HiFi reads and normalizes their cumulative length using X-degenerate-region-specific k-mers.

Nucleotide-resolution characterization of these regions has been similarly challenging because of their highly repetitive structure. Yq12 is almost entirely absent from the human reference genome (GRCh38; Figure 1a)^2^. The combination of Oxford Nanopore ultra-long and PacBio HiFi sequencing has enabled complete nucleotide-level resolution of Yq12 in 84 of these 142 Y chromosomes despite the exceptional complexity of this region^6^. These assemblies revealed extensive interindividual variation in Yq12 length, ranging from approximately 10.3 to 53.1 Mb, with a mean of 27.1 Mb^6^. Despite extensive variation in total Yq12 length and repeat-array organization among individuals, analyses of completely assembled Yq12 regions have shown that *DYZ1* and *DYZ2* copy numbers tend to remain close to a 1:1 ratio across males^5,6^. The persistence of this near-1:1 copy-number ratio across males is suggestive of coordinated changes in *DYZ1* and *DYZ2* abundance. However, cross-sectional comparisons alone may not be sufficient to establish whether the near-1:1 *DYZ1*:*DYZ2* ratio reflects a constrained evolutionary state maintained over time.

Substantial interindividual variation in Yq12 heterochromatin size and *DYZ1*/*DYZ2* repeat content has long been documented among phenotypically normal males^16–19^. Differences in Yq12 repeat content or organization have also been reported among unrelated individuals^8,20^, monozygotic twins^21^, and fathers and sons^22^, consistent with a high rate of change in this region, although these earlier studies did not provide nucleotide-resolved or population-scale estimates of total Yq12 length. Complete pedigree assemblies further indicate elevated sequence turnover within Yq12, which shows a higher per-nucleotide *de novo* mutation rate than euchromatic regions of the Y chromosome, with two pedigree analyses reporting averages of 9.6^23^ and 10.8^23^ *de novo* germline single-nucleotide variants per transmission.

At the same time, extreme Yq12 repeat content and structural rearrangements have been associated with reproductive abnormalities, including recurrent pregnancy loss, impaired embryo development, and male infertility^24–27^. An earlier attempt to quantify this variation using Illumina whole-genome sequencing (WGS) from 396 males in the 1000 Genomes Project (1kGP) suggested a total combined *DYZ1* repeat length variation between 7 and 98 Mb, with a mean of 24 Mb^8^. The upper end of this estimated range (98 Mb) is approximately 45 Mb larger than the largest Yq12 region observed among the 84 completely assembled Y chromosomes (53.1 Mb)^6^, underscoring the need for an assembly-benchmarked method. Genome assemblies now indicate that *DYZ1* constitutes approximately 60% of Yq12 by sequence length^5–7^, consistent with a near-1:1 copy-number ratio given the larger *DYZ1* repeat unit (≈3.5 kb versus ≈2.4 kb). Complete assemblies of the Yq12 region have transformed our understanding of Yq12 structure and composition, but they remain too few to characterize variation at population scale.

Although satellite arrays are frequently associated with constitutive heterochromatin, transcription of satellite-derived RNAs is well documented in multiple species^28–31^. These transcripts have been implicated in the establishment and maintenance of heterochromatin and in higher-order nuclear organization^32,33^. Evidence for transcription from Yq12 remains limited, but includes at least two distinct polyadenylated *DYZ1*-derived noncoding RNAs with testis-specific expression, one of which contributes Yq12-derived sequence to a testis-specific chimeric *CDC2L2* transcript^34^. More recently, HSat3 transcription from Yq12 has also been observed in cultured male cells following heat shock, where the locus recruits HSF1 and contributes to the formation of nuclear stress bodies^35^. Because *DYZ1* is a Y-linked HSat3 variant, and large HSat3 arrays have recently been shown to act as megabase-scale transcription-factor-binding platforms in cultured cells^36^, Yq12 transcription is of particular interest. However, the cell types responsible for Yq12 transcription within the human testis and its dynamics across spermatogenesis remain poorly understood.

To investigate the genetic variation, evolutionary dynamics, and transcriptional activity of Yq12, we developed SRKmer and LRKmer, k-mer-based methods that estimate Yq12 length directly from short- and long-read whole-genome sequencing data without requiring a sample-specific assembly and with reliable performance at low sequencing depth. We applied these methods to characterize population- and haplogroup-associated variation in Yq12 length, test whether Yq12 length and *DYZ1*:*DYZ2* repeat composition are evolutionarily constrained, compare Yq12 estimates in father-son pairs, and examine Yq12 repeat expression across human testis transcriptomic datasets. Together, this work provides a scalable framework for investigating the structural variation, evolutionary behavior, and transcriptional activity of one of the largest heterochromatic regions of the human genome.

## Materials and Methods

### Short-read K-mer Estimator

The short-read k-mer estimator (SRKmer: https://github.com/Markloftus/Yq12Estimations/tree/main/SRKmer) was written in Python/Cython to utilize Illumina short-read WGS data to estimate Yq12 length, as well as *DYZ1* and *DYZ2* repeat copy numbers (note: only short-read data generated using a PCR-free library preparation were found to produce accurate estimates of the Yq12 region, data not shown). The SRKmer was designed to run on BAM/CRAM files read using Pysam (v0.22.0) (https://github.com/pysam-developers/pysam)^37–39^. A Yq12 repeat k-mer library was constructed from a set of all 24-base pair long k-mers derived from 7 previously assembled Yq12 regions^5^. A k-mer length of 24 bp was selected to maximize sequence specificity while maintaining a computationally manageable library size. Each k-mer was screened against the CHM13v2 human reference genome to determine whether it occurred outside the Yq12 region; any k-mer with one or more matches elsewhere in the genome was removed. SRKmer also uses a second 24-base k-mer library derived from satellite sequences in the T2T-CHM13v2 Y-chromosome centromere to identify and filter centromeric-derived reads, thereby reducing their potential contribution to the Yq12 estimate.

The sequence of each read in the BAM/CRAM file was queried against these libraries of Yq12-derived and centromeric-derived 24-mers. Reads were retained only if they contained at least one Yq12-derived 24-mer for which all 24 bases had base quality ≥20. The SRKmer uses sequencing coverage across the X-degenerate regions (XDR) of the Y chromosome to normalize for read depth. The XDR coordinates (**Supplementary Table S1)** were divided into 1-kb windows, and coverage within each window was calculated using Samtools (v1.15.1)^38^ (samtools bedcov −Q 20). Mean coverage across all XDR windows was used as the estimated Y chromosome sequencing depth. The sample Yq12 length (*L*) is calculated as the sum of all read lengths (*H*) divided by the average XDR-derived read depth (*D*):

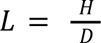

SRKmer produces two estimates of Yq12 size. The first is based on all reads classified as Yq12-derived using the full k-mer library: the summed length of these reads is divided by the mean XDR coverage to estimate the total length of Yq12. Yq12-derived reads used for the second estimate were identified based on the presence of k-mers with ≥95% sequence identity to the *DYZ1* and *DYZ2* repeat consensus sequences, with sequence identity determined using Hamming distance^40^. The summed length of these reads is similarly divided by the mean XDR coverage. The second approach also enables estimation of *DYZ1* and *DYZ2* repeat-unit copy numbers. Reads were assigned to repeat-subunit-specific bins, and their lengths are summed within each bin. The resulting length estimates are then divided by the average subunit length (3569 bp for *DYZ1* and 2420 bp for *DYZ2*) to estimate subunit copy number.

### Long-read K-mer Estimator

The long-read k-mer Yq12 estimation method (LRKmer: https://github.com/Markloftus/Yq12Estimations/tree/main/LRKmer) was written in Python and utilizes Yq12-derived k-mers to calculate the percentage of a HiFi read that is composed of Yq12-derived sequence. The LRKmer was designed to run on HiFi-based gzip FASTQ files (FASTQ.gz). Similarly, a Yq12-derived k-mer library was constructed from 84 males with previously assembled, complete Yq12 regions^6^. Yq12 sequences were retrieved using Samtools/Pysam (v0.22.0) (https://github.com/pysam-developers/pysam)^37–39^ and all 24-mers were extracted. K-mers occurring only once were excluded to minimize spurious signals and reduce the overall size of the k-mer library. Unlike SRKmer, k-mers used by LRKmer were permitted to occur outside Yq12 because LRKmer scans long sequencing reads for multiple Yq12-associated k-mers, rather than inferring Yq12 origin from a single k-mer match as in short-read data. In a benchmark subset of 74 previously assembled Yq12 regions^6^ (i.e., 74 of the 84 complete Yq12 assemblies were from open-access samples that overlapped the 1kGP dataset), estimated Yq12 length decreased as the minimum proportion of Yq12-derived k-mers required per HiFi read increased, but largely plateaued at thresholds of 50% and above (**Supplementary Figure S1**). Additionally, the average percent difference between k-mer-based estimates of Yq12 lengths and the *de novo* assembled Yq12 lengths was lowest (i.e., most accurate; mean: 3% difference) when reads that contain 50% or more Yq12-derived k-mer sequence are retained in the calculation **(Supplementary Figure S2)**. As such, all reads whose sequence contained 50% or more Yq12-derived k-mers were retained for all LRKmer-based Yq12 length estimations. This resulted in a final library of 4,308,723 24-base pair long Yq12-derived k-mers. Similar to the SRKmer, read depth across the XDRs is taken into account for the Yq12 length estimations. The sequence for each XDR within the CHM13v2 Y reference chromosome (chrY:2458321-2726838, chrY:6400614-7238446, chrY:12656254-14891093, chrY:14964945-16781318, chrY:16811411-17066091, chrY:17332353-18362321, chrY:19776674-20961373, chrY:21226252-22211007) was retrieved using Samtools/Pysam (v0.22.0) (https://github.com/pysam-developers/pysam)^37–39^ and all 24-mers occurring exactly once were retained. This resulted in a library of 6,430,092 unique XDR-derived k-mers. The LRKmer keeps track of how many reads contain these unique XDR-derived k-mers. The average count of total reads that contain these k-mers across all XDRs is utilized as the putative sample read depth.

### Read-depth subsampling and whole-genome simulation

To evaluate the effects of sequencing depth on Yq12 length estimation, publicly available Illumina short-read and PacBio HiFi whole-genome sequencing data for HG01890 were obtained from the Human Genome Structural Variation Consortium (HGSVC)^7^. HG01890 was randomly selected from the group of 84 samples which have a previously assembled Yq12 region^6^, allowing estimated lengths to be compared with the assembled Yq12 length. The short-read dataset, with an original Y-chromosome sequencing depth of approximately 15.5x, was randomly subsampled to 10%, 25%, and 50% of the original read count, corresponding to approximate Y-chromosome depths of 1.6x, 3.9x, and 7.8x, respectively. The HiFi dataset utilized, with an original Y-chromosome depth of approximately 6x, was randomly subsampled to 16%, 33.3%, and 50% of the original reads, corresponding to approximate Y-chromosome depths of 1x, 2x, and 3x, respectively. Five independent random subsamples were generated at each depth. Estimation accuracy was defined as 100 − (|estimated − assembled| / assembled x 100), such that 97% accuracy corresponds to an estimate within 3% of the assembled Yq12 length. Reported values represent the lowest accuracy observed across all five replicates at a given depth.

SRKmer and LRKmer were applied to the short- and long-read subsamples, respectively, using the same parameters applied to the original datasets, and the resulting estimates were compared with the assembled HG01890 Yq12 length. To further evaluate LRKmer using controlled (sequence bias free) whole-genome data, error-free long reads were simulated from the CHM13v2.0 reference genome using custom scripts implemented with Pysam (v0.22.0)^37–39^ and Biopython (v1.82)^41^. For each chromosome, read start positions were selected randomly, and read lengths were sampled between 100 and 30,000 bp. Reads were generated until the cumulative read length reached the specified target coverage and were written as compressed FASTQ files. The simulation procedure was applied at target whole-genome coverages ranging from 1x to 60x (1x, 2x, 4x, 8x, 15x, 30x and 60x). Each simulated dataset was analyzed with LRKmer using the same parameters applied to real datasets.

### Male Sample Selection For Yq12 Estimation

In total, 1,597 male human genomes from the expanded 1kGP cohort were analyzed^42,43^. Short-read Illumina whole-genome sequencing data of 1,598 males, including 309 father-son pairs, were retrieved from the high-coverage 1kGP dataset (sequenced to an average depth of approximately 30x)^42^. One male sample, NA19226, was excluded because its BAM/CRAM file consistently produced errors during processing, leaving 1,597 males for SRKmer analysis. For the LRKmer-based Yq12 estimation, long-read PacBio HiFi data were obtained from the HGSVC (31 males)^7^ and the Human Pangenome Reference Consortium (HPRC; 114 males; https://data.humanpangenome.org/raw-sequencing-data)^44^. Sample ID information for SRKmer and LRKmer analyses is provided in **Supplementary Tables S2 and S3**.

### Y haplogroup determination

Joint genotyping of the 1kGP Illumina WGS data^42^ was performed as described previously^5^. Analyses were restricted to approximately 10.4 Mb of chromosome Y previously defined as accessible to short-read sequencing^45^. Variants were called using BCFtools (v1.9)^38,46^, assuming haploid ploidy and requiring minimum base and mapping qualities of 20. Indels were removed, as were SNVs located within 5 bp of an indel call using the SnpGap filter. Genotype calls were further required to have a minimum read depth of three. At positions supported by more than one allele, the called allele was required to account for at least 85% of the reads; calls not meeting this threshold were set to missing. Seven samples (HG02966, HG03008, HG01982, HG01984, HG03998, HG03006, HG03976) with >15% missing genotypes were excluded, as were sites with ≥5% missing genotypes, using VCFtools (v0.1.16)^47^, leaving a total of 1,590 samples. Y-chromosome haplogroups were assigned using yHaplo software (https://github.com/23andMe/yhaplo)^48^, as previously described^49^, using nomenclature from version 15.73 of the International Society of Genetic Genealogy (ISOGG) Y-DNA haplogroup tree. A maximum-likelihood tree was inferred with RAxML (v8.2.10)^50^ under a GTR plus gamma model using filtered chrY variant sites.

### Population-scale phylogenetic analysis of Yq12 size and composition

Phylogenetic analyses were performed using SRKmer estimates and the maximum-likelihood Y-chromosome phylogeny (see **Y haplogroup determination methods**), with 1,590 males represented in both datasets. The primary tree was rooted using HG01890 as the outgroup, with the root placed at the midpoint of the branch leading to HG01890. *DYZ1* and *DYZ2* repeat-subunit counts were log2-transformed, repeat composition was represented as log2(*DYZ1*/*DYZ2*), such that parity corresponded to zero, and total Yq12 length was analyzed as log2-transformed base-pair length. For evolutionary model fitting, branch lengths were scaled so that the mean root-to-tip distance equaled one to improve numerical stability. Models included independent tip-level residual variance. Phylogenetic signal in *DYZ1*:*DYZ2* composition was quantified using Pagel’s λ^51,52^, and evolutionary coupling between *DYZ1* and *DYZ2* was evaluated using a bivariate phylogenetic variance-covariance model and simulations under a zero-covariance null. From the fitted covariance matrix, the evolutionary correlation and variance along shared-abundance and *DYZ1*:*DYZ2* imbalance axes were calculated; bootstrap confidence intervals and Monte Carlo tests were used to evaluate these statistics and differences in marginal evolutionary variance between *DYZ1* and *DYZ2*. Phylogenetic generalized least-squares regression^51^ was utilized to test the relationship between log2 Yq12 length and *DYZ1*:*DYZ2* composition.

To test whether total Yq12 length and *DYZ1*:*DYZ2* composition were evolutionarily constrained, traits were modeled under Brownian-motion and Ornstein–Uhlenbeck (OU) models^53,54^. For Yq12 length, Brownian motion was compared with both a nonstationary OU model with freely estimated optimum and root state and a stationary OU model with a freely estimated optimum. In the stationary model, the ancestral state was integrated over the stationary OU distribution rather than estimated as an independent root value. Model support was evaluated using AIC^55^, and significance of the OU advantage over Brownian motion was assessed by parametric bootstrap in which datasets were simulated under the fitted Brownian-motion model, and all three models were refitted to each simulated dataset. Of 1,000 bootstrap datasets generated, 995 yielded successful fits for all three models and were retained for analysis. As a complementary descriptive test, the width containing the central 90% of observed Yq12 lengths was compared with its distribution under the Brownian-motion simulations. Model fitting was repeated after midpoint rooting to evaluate sensitivity to root placement (see **code availability**).

For *DYZ1*:*DYZ2* composition, Brownian motion was compared with stationary OU models in which the compositional optimum was either fixed at parity or estimated freely. Support for the parity-centered stationary OU model over Brownian motion was evaluated using AIC^55^, with significance assessed from 500 datasets simulated under the fitted Brownian-motion model and refitted under both models. As a complementary consistency check, latent ancestral states were reconstructed, and both a branch-length-weighted restoring slope and the fraction of branches moving toward parity were calculated, with significance assessed against 20,000 Brownian-motion simulations subjected to the same reconstruction procedure (see **code availability**).

Major Y haplogroups were additionally tested for distinct compositional optima using a multi-regime stationary OU model^53^. Seven monophyletic haplogroups represented by at least 40 males (E1, R1, O1, J2, Q1, H1, and R2) were assigned separate optima and compared with the single-optimum OU model using AIC, BIC, likelihood-ratio testing, and a 500-replicate parametric bootstrap. In the primary model, each regime included the branch entering the haplogroup most recent common ancestor and its descendants. Sensitivity analyses repeated the OU models under midpoint rooting, placed lineage-specific shifts at the crown node while excluding the entering stem branch, and repeated the multi-regime comparison after subtracting the fitted phylogenetic generalized least-squares regression effect of log2 Yq12 length from log2(*DYZ1*/*DYZ2*).

### Human Testis Data

Previously generated, publicly available PacBio Iso-Seq and single-cell RNA-sequencing (scRNA-seq) data from human testis were analyzed. The PacBio Iso-Seq data (SRR12544673 and SRR12544672) were generated from pooled normal human testis samples and retrieved from BioProject PRJNA659539^56^. The scRNA-seq was originally performed using the 10x Genomics system (paired-end Illumina HiSeq 2500) on three healthy male donors (donor 1 (D1): 17 years of age; donor 2 (D2): 24 years of age; donor 3 (D3): 25 years of age). The scRNA-seq data (SRR6860519 - D1/rep1, SRR6860520 - D1/rep2, SRR6860521 - D2/rep1, SRR6860522 - D2/rep2, SRR6860523 - D3/rep1, SRR6860524 - D3/rep2) were retrieved from accession GEO: GSE120508 and GSE112013^57^.

### Testis scRNA-seq and Iso-Seq Analyses

All scRNA-seq reads^57^ were screened for Yq12 repeat (*DYZ1* and *DYZ2*) expression using the SRKmer estimator. Only repeat k-mers assigned to *DYZ1* or *DYZ2* with a classification similarity greater than 0.95 were retained. For each Yq12-positive read, the corresponding BAM record was used to recover the corrected 10x Genomics cell barcode (CB) and unique molecular identifier (UB; the raw UMI tag UR was used when UB was unavailable). Secondary and supplementary alignments were excluded, and each sequencing-read name was processed only once. Cell barcodes were matched to the published cell metadata to assign both broad and refined cell-type annotations. Reads with conflicting *DYZ1* and *DYZ2* classifications, as well as cell barcode/UMI combinations supported by both repeat classes, were excluded as ambiguous. To remove PCR amplification duplicates, reads were collapsed by the unique combination of sequencing library, cell barcode, UMI, and repeat class. Thus, multiple reads or k-mer matches carrying the same cell barcode, UMI, and repeat classification were counted as one repeat-containing molecule. *DYZ1* and *DYZ2* abundance was calculated separately for each cell type and technical replicate. Repeat-specific UMI counts were divided by the total number of UMIs captured from all cells assigned to that cell type and multiplied by 10,000. Additionally, scRNA-seq reads were screened for satellite composition using RepeatMasker (v4.1.7)^58^ (-s slow parameter) and the Dfam (v3.8) library^59,60^. Testis Iso-Seq reads were screened for Yq12 repeat (*DYZ1* and *DYZ2*) expression using the LRKmer estimator. Iso-Seq reads composed of ≥90% *DYZ1* and *DYZ2* k-mer sequences were retrieved using Samtools (v1.15.1)^38,39^, aligned and visualized against the *DYZ1* and *DYZ2* consensus sequences using AliView (v1.30)^61^.

### Father-Son Comparisons

Father-son analyses initially included 309 pairs. Six pairs with differences of at least 20% in estimated Yq12 length were excluded from the primary statistical analysis, leaving 303 pairs. Father-son differences in estimated Yq12 length were evaluated using a two-sided Wilcoxon signed-rank test, and the numbers of pairs in which the father or son had the longer estimated Yq12 were compared using a two-sided exact binomial sign test; both tests were implemented in SciPy (v1.11.1)^62^. To determine whether father-son Yq12 length differences varied among Y-chromosome haplogroups, a Kruskal-Wallis test was performed using SciPy for haplogroups represented by at least 10 father-son pairs. Two-sided Fisher’s exact tests (as implemented in SciPy) were used to evaluate associations between whether the father’s Yq12 length was above or below the global or haplogroup-specific median and whether the son’s estimated Yq12 was longer or shorter than the father’s. Conditional maximum-likelihood odds ratios with exact 95% confidence intervals were computed with SciPy (scipy.stats.contingency.odds_ratio). For father-son *DYZ1*:*DYZ2* comparisons, subunit ratios were natural-log transformed, such that a balanced ratio of 1 corresponded to zero. The proportion of pairs in which the son’s ratio was closer to 1 than the father’s was evaluated using a one-sided exact binomial sign test in SciPy, with the alternative hypothesis that the proportion exceeded 0.5. A 95% Clopper-Pearson exact confidence interval for this proportion was calculated using statsmodels.

### Statistical Analyses

All statistical analyses were performed in Python (v3.11.5) using NumPy (v1.24.3)^63^, pandas (v2.0.3)^64,65^, SciPy (v1.11.1)^62^, statsmodels (v0.14.0)^66^, and scikit-posthocs (v0.7)^67^. Pearson correlation coefficients were calculated using SciPy to evaluate linear relationships between estimated and assembled Yq12 lengths, between SRKmer and LRKmer estimates, between assembled and SRKmer-estimated *DYZ1*:*DYZ2* subunit ratios, and between estimated *DYZ1* and *DYZ2* subunit counts. Rank concordance between estimated and assembled Yq12 lengths was evaluated using Spearman’s rank correlation and Kendall’s τ^68^ in SciPy. Differences in estimated Yq12 length among Y-chromosome haplogroups were evaluated using the Kruskal-Wallis test in SciPy, restricting the analysis to haplogroups represented by more than 10 males. Kruskal-Wallis epsilon-squared was calculated as an effect-size estimate. Pairwise haplogroup comparisons were performed using Dunn’s tests in scikit-posthocs with Benjamini-Hochberg false-discovery-rate correction^69^. Differences between haplogroups were further summarized using group medians and Hodges-Lehmann location-shift estimates with 95% confidence intervals obtained from 3,000 bootstrap resamples using custom functions implemented with NumPy and pandas.

## Results

### SRKmer and LRKmer estimates are strongly correlated with assembled Yq12 lengths

Population-scale characterization of Yq12 variation requires accurate approaches for estimating its length and repeat composition without relying on complete Y-chromosome assemblies. To address this need, we developed two independent k-mer-based approaches: the SRKmer and LRKmer estimators (**Methods**). Both approaches use 24-mers from high-quality, fully assembled Yq12 regions^5,6^ to identify Yq12-specific reads. Yq12 length is then estimated by summing the identified sequence and normalizing using XDR-based coverage estimates (**Figure 1b,c**; **Methods**)^5,6^. Next, we assessed the accuracy of both estimators across 74 samples from the 1kGP (i.e., overlapping benchmark set) that were also among the 84 samples with complete, high-quality Yq12 assemblies^6^. Estimates from both methods correlated strongly with assembled Yq12 lengths (SRKmer: Pearson’s r = 0.986, p-value=2.68e^−57^; LRKmer: Pearson’s r = 0.869, p-value=1.01e^−23^; **Figure 2a; Supplementary Tables S2 and S3**). To assess whether the two estimators captured consistent inter-individual variation in Yq12 length, we compared SRKmer and LRKmer estimates across the same 74 samples. The estimates were strongly correlated (Pearson’s r = 0.87, p-value=3.10e⁻²⁴; **Supplementary Figure S3**). SRKmer additionally estimates repeat-specific *DYZ1*:*DYZ2* copy-number ratios, a capability not implemented in LRKmer. SRKmer-estimated ratios correlated strongly with assembly-derived ratios (Pearson’s r = 0.952, p-value=2.53e⁻⁴⁰; **Figure 2b**). Rank-based comparisons further assessed whether each estimator preserved the relative ordering of samples by Yq12 length. SRKmer showed strong concordance between estimated and assembled Yq12 length ranks (Spearman’s ρ = 0.979, p-value=7.93e⁻⁵²; Kendall’s τ = 0.886, p-value=5.88e⁻²⁹), while LRKmer also showed strong rank concordance (Spearman’s ρ = 0.817, p-value=6.50e⁻¹⁹; Kendall’s τ = 0.648, p-value=3.05e^−16^). Preservation of relative sample ordering provides additional confidence that comparisons of Yq12 length among samples are robust to residual estimation error.

**Figure 2:**
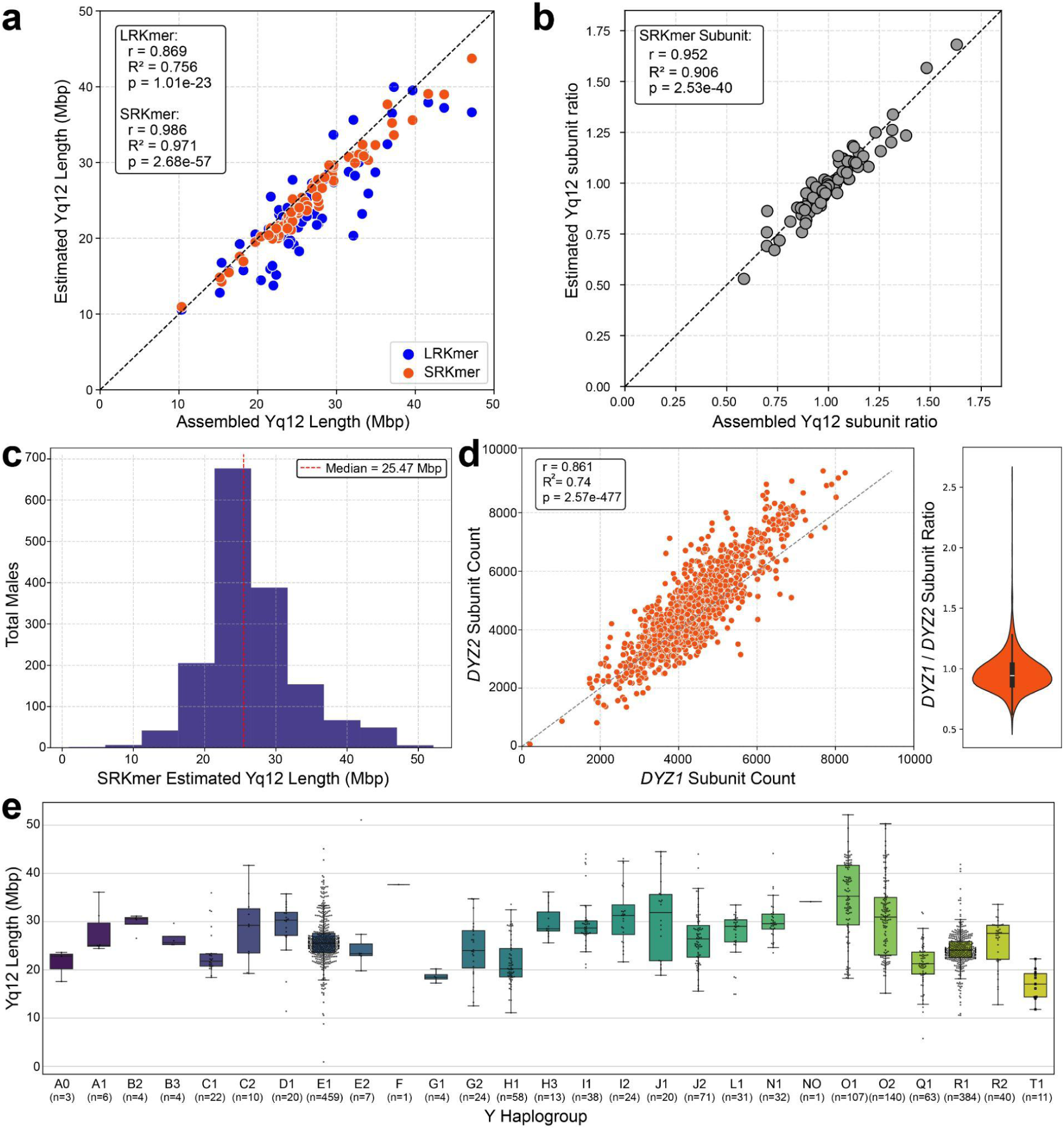
Validation and population-scale application of Yq12 estimators. **a**) Comparison of SRKmer- and LRKmer-estimated Yq12 lengths with lengths measured directly from *de novo* assemblies across 74 males. **b**) Comparison of assembly-derived and SRKmer-estimated *DYZ1:DYZ2* repeat ratios for the same samples. Each point represents one male; the dashed black line indicates perfect accuracy (y = x). **c**) Distribution of SRKmer-estimated Yq12 lengths across 1,597 males. The dotted red line indicates the median. **d**) Relationship between estimated *DYZ1* and *DYZ2* repeat copy numbers (left) and distribution of *DYZ1:DYZ2* repeat ratios (right). Slightly lower *DYZ1* estimates may reflect variation in *DYZ1* repeat-unit length^5^, as copy number was calculated using a fixed unit length of 3,569 bp. e) Distribution of SRKmer-estimated Yq12 lengths across major Y-chromosome haplogroups. Boxes show the median and interquartile range.

Both estimators remained accurate at reduced sequencing depths, with consistent Yq12 length estimates across independent subsampling replicates. In HG01890, a benchmark sample with a previously assembled Yq12 region and both short- and long-read WGS data, SRKmer maintained >97% accuracy across the tested short-read depths (1.6-7.8x), while LRKmer maintained >88% accuracy across the tested long-read depths (1-3x) **(Supplementary Figure S4a,b; Methods)**. **In bias-free simulated whole-genome HiFi data, LRKmer achieved accuracy comparable to SRKmer**, with estimates within 2.5% of the CHM13v2 Yq12 length across all tested coverage levels from 1x to 60x and within 1% at five of seven coverage levels **(Supplementary Figure S4c)**. These results indicate that both estimators remain stable at substantially reduced sequencing depths. Given its higher accuracy in our benchmarking and the availability of Illumina short-read data for 1,597 males in the 1kGP, we used SRKmer for all subsequent Yq12 length analyses.

### Population-scale estimates reveal extensive Yq12 length variation across 1,597 males

To determine the size variation and evolution of the Yq12 region at the population level and across a wide range of haplogroups, we ran the SRKmer tool on PCR-free, high-coverage Illumina WGS data from 1,597 males from the 1kGP cohort^42^. The estimated Yq12 length ranged from 0.93 to 52.1 Mb (Yq12 length mean: 26.5 Mb, median: 25.5 Mb) (**Figure 2c; Supplementary Table S2**). The majority of males (67.25%; n = 1,074) had a Yq12 length between 20 and 30 Mb. In contrast, only 6 males (0.37%) had estimates below 10 Mb or above 50 Mb. These six had chrY sequencing depths of 15.8-20.0x, spanning the cohort median (16.1x) and with none falling below the cohort first quartile (15.4x), indicating that low coverage does not account for their extreme length estimates. Across the overall cohort, *DYZ1* and *DYZ2* subunit counts remained strongly correlated even though total Yq12 length varied more than 50-fold (Pearson’s r = 0.861, p-value=2.57e^−477^). The estimated *DYZ1:DYZ2* ratio had a median of 0.944 and a mean of 0.963, with values ranging from 0.529 to 2.59 (**Figure 2d; Supplementary Table S2**). The broad population distribution of *DYZ1*:*DYZ2* ratios was centered near parity, consistent with the repeat composition observed in complete assemblies^5,6^.

We next examined the relationship between estimated Yq12 length and Y-chromosome haplogroup. The 1,597 males represented 27 major Y haplogroups, and Yq12 length varied substantially both within and among haplogroups (**Figure 2e; Supplementary Tables S2 and S3**). To compare Yq12 length distributions among haplogroups, we restricted the analysis to the 18 haplogroups represented by more than 10 males (11-459 males per haplogroup). Yq12 length differed significantly among these groups (Kruskal-Wallis: H = 407.84, p-value=4.25e⁻⁷⁶; ε² = 0.254), corresponding to a large rank-based effect of haplogroup membership. Post hoc Dunn’s tests with Benjamini–Hochberg false-discovery-rate correction identified multiple significant pairwise differences, while per-group medians and Hodges–Lehmann differences with 95% confidence intervals quantified the magnitude of these contrasts (**Supplementary Tables S4–S7**). Among the sampled haplogroups, O1 had the largest median Yq12 estimate (35.26 Mb), whereas T1 had the smallest (17.08 Mb).

We next asked whether Yq12 length could be explained by unconstrained evolutionary change or instead showed evidence of long-term constraint. To test this, we modeled log2-transformed Yq12 length across the Y-chromosome phylogeny using a Brownian-motion model, in which evolutionary variance accumulates without a preferred size, and a stationary Ornstein–Uhlenbeck (OU) model, in which values tend to remain around an inferred equilibrium. The stationary OU model fit the data substantially better than Brownian motion (ΔAIC = 61.26; parametric-bootstrap p-value=0.0010; **Supplementary Table S8**), supporting long-term constraint on Yq12 length. The inferred equilibrium was 26.24 Mb, with an approximate central 95% stationary range of 15.8–43.6 Mb, indicating constraint around a broad range of Yq12 sizes. This result was robust to midpoint rooting, which produced an essentially identical equilibrium estimate (26.24 Mb) and similarly strong support for OU over Brownian motion (ΔAIC = 61.80; **Supplementary Table S8**).

### *DYZ1*:*DYZ2* composition is evolutionarily constrained near parity

Next, we examined repeat composition in the context of the Y-chromosome phylogeny across 1,590 samples (**Methods**) because the maintenance of a roughly 1:1 *DYZ1*:*DYZ2* ratio observed across males could partly reflect shared ancestry. *DYZ1*:*DYZ2* composition showed a strong phylogenetic signal (Pagel’s λ = 0.990), indicating that closely related Y chromosomes tended to have similar repeat compositions. Despite this lineage structure, *DYZ1* and *DYZ2* abundance changed largely in concert across the phylogeny (ρ = 0.855; 95% bootstrap CI, 0.842–0.868). Consistent with this, evolutionary variance along the axis representing shared changes in *DYZ1* and *DYZ2* abundance was 10.33-fold greater than variance in their relative imbalance (100,000-replicate Monte Carlo test under a zero-covariance null, p-value=1.0e^−5^). Thus, most evolutionary change appeared to involve joined expansion or contraction of both repeat families rather than large shifts in their relative proportions. However, the two repeat families were not completely symmetric: *DYZ2* showed 1.74-fold greater evolutionary variance than *DYZ1* (Monte Carlo test under an equal-variance null, p-value = 1.0e^−5^), and an increase in Yq12 length was associated with a lower *DYZ1*:*DYZ2* ratio in phylogenetic generalized least-squares regression (slope = −0.280; 95% CI, −0.308 to −0.253; p-value=1.98e^−7^^8^; **Supplementary Table S9**). Together, these results indicate that Yq12 expansion and contraction generally involve coordinated changes in both repeat families, with *DYZ2* contributing disproportionately to variation in larger Yq12 arrays.

Given the broad clustering of *DYZ1*:*DYZ2* ratios near parity across males despite substantial variation in Yq12 size, we next asked whether this compositional pattern reflected unconstrained evolutionary drift or long-term constraint around a characteristic ratio. A stationary Ornstein–Uhlenbeck model centered at parity fit the data substantially better than a Brownian-motion model of unconstrained drift (ΔAIC = 13.37; 500-replicate parametric bootstrap under the fitted Brownian-motion null, p-value=0.002). This result supports long-term constraint of *DYZ1*:*DYZ2* composition around an approximately 1:1 repeat ratio. When we estimated the equilibrium ratio directly from the data, the optimum was 0.932 and did not significantly improve model fit relative to a fixed ratio of 1.0 (likelihood-ratio test, p-value=0.236; **Supplementary Table S10**), indicating that the inferred equilibrium is close to parity. We also found no evidence that different Y-chromosome lineages maintain different preferred ratios: allowing seven major Y haplogroups to have separate compositional optima did not improve model fit (500-replicate parametric-bootstrap likelihood-ratio test under the single-optimum OU model, p-value=0.948; **Supplementary Table S11**). Finally, individual evolutionary changes showed no detectable bias toward restoring parity (48.0% moved toward parity; Monte Carlo test against 20,000 Brownian-motion simulations, *P* = 0.722; **Supplementary Table S10**). Thus, the near-parity pattern appears to reflect constraint operating over longer evolutionary timescales rather than systematic correction toward parity after each individual change.

### Exploratory father-son comparisons show asymmetric paired estimates

Three father-son transmissions with complete Yq12 assemblies^6^ were available from a previously analyzed four-generation CEPH pedigree^23^: NA12877-NA12886, NA12877-NA12882, and 200080-200084. In two of the three pairs, the paternal Yq12 assembly was longer than the son’s by 839 bp and 30,401 bp, respectively, whereas in the third pair, the son’s Yq12 was longer by 29,223 bp^6^. We extended this analysis to 309 father-son pairs from our 1kGP set of 1,597 males (**Supplementary Table S12; Figure 3a**) using the SRKmer estimator to evaluate size and the direction of intergenerational differences. For the primary statistical analysis, we excluded six father-son pairs with ≥20% length differences in estimated Yq12 length to reduce the potential influence of large deletion or duplication events that may not reflect typical intergenerational variation, leaving 303 pairs. Across the 303 father-son pairs, sons had, on average, significantly shorter estimated Yq12 regions than their fathers, with fathers carrying the longer estimated Yq12 in 214 pairs (70.6%) (Wilcoxon signed-rank test W=11,089, two-sided p-value=5.20e^−15^; two-sided exact binomial sign test: p-value = 4.92e^−13^, **Figure 3b; Supplementary Figure S5**). To assess whether exclusion of the six father-son pairs with differences of at least 20% influenced the results, we repeated the analysis using all 309 pairs, which yielded consistent results (in 219 pairs the father’s Yq12 was larger; Wilcoxon signed-rank test: W=11,400, two-sided p-value=1.43e^−15^; two-sided exact binomial sign test: p-value=1.46e^−13^).

**Figure 3:**
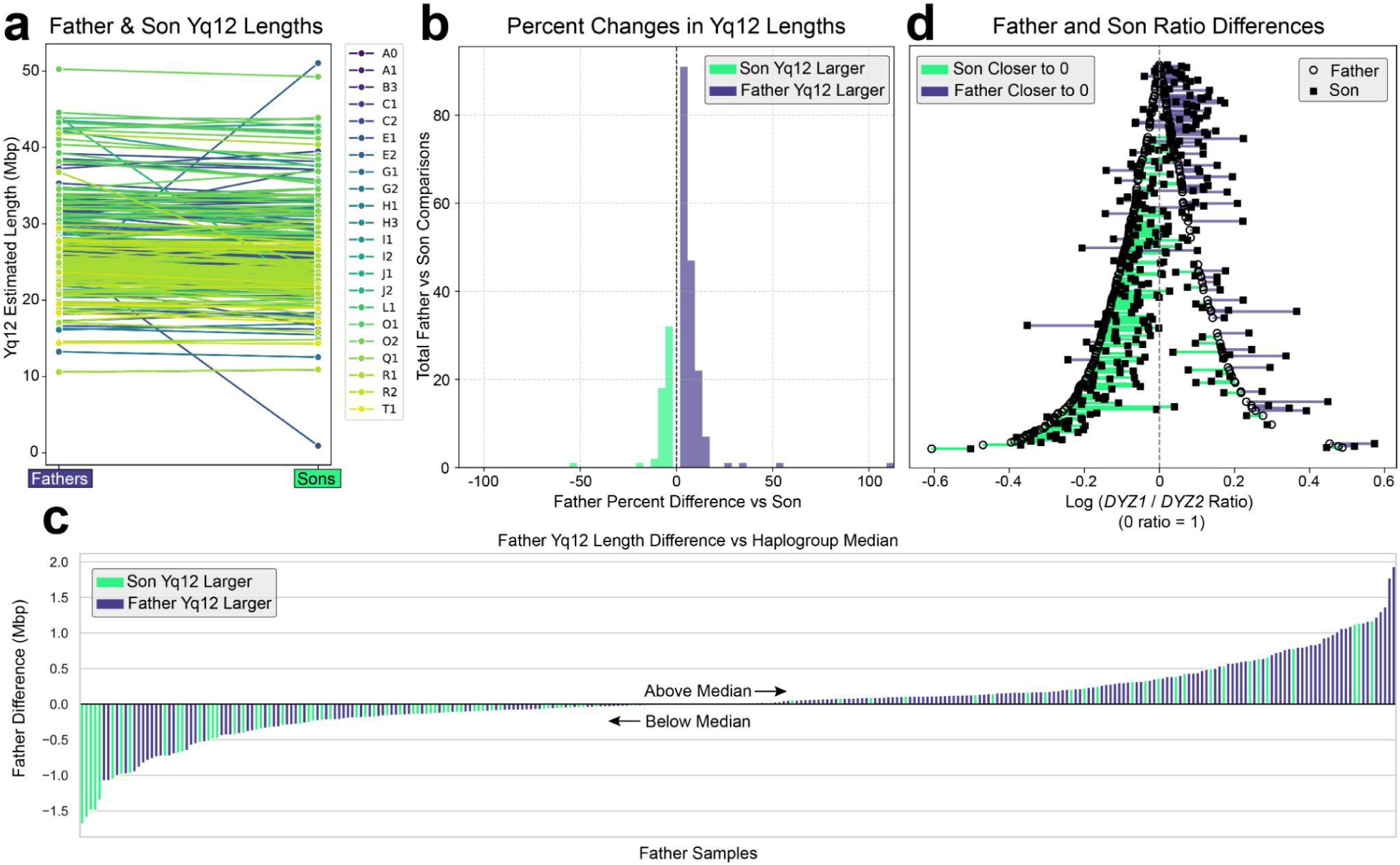
Father-son variation in Yq12 length and repeat composition. **a**) SRKmer-estimated Yq12 lengths in 309 father-son pairs, with paired values connected and colored by major Y haplogroup. **b**) Distribution of percentage differences in Yq12 length in the son relative to the father (309 pairs). Purple indicates a longer Yq12 in the father and seafoam green a longer Yq12 in the son. **c**) Difference between each father’s estimated Yq12 length and the median for his Y haplogroup, ordered from the largest negative to largest positive deviation. Bars are colored according to whether Yq12 is longer in the son (seafoam green) or father (purple) in 303 father-son pairs (excluding six pairs with Yq12 length differences ≥20%). **d**) Log-transformed SRKmer-estimated *DYZ1*:*DYZ2* repeat ratios in 303 father-son pairs. Fathers are shown as circles and sons as squares; lines indicate whether the son’s ratio is closer to parity (i.e., 0, seafoam green) or further away from parity (purple) than the father’s.

Our findings and previous studies^6,8^ indicate variation in Yq12 length and repeat architecture among Y haplogroups, suggesting that lineage-specific Yq12 organization could shape patterns of intergenerational variation. To test this, we determined father-son variation stratified by haplogroup. Requiring at least 10 pairs per given haplogroup, we retained 83.5% of pairs (253/303) across five haplogroups (E1: 121 pairs, R1: 76 pairs, O2: 28 pairs, Q1: 15 pairs, and O1: 13 pairs). We found no evidence that father-son differences vary by haplogroup (Kruskal-Wallis: H = 3.55, df = 4, p-value=0.471), though with 253 pairs across five haplogroups the analysis has limited power to detect modest differences.

We next investigated whether the direction of father-son length differences was associated with the father’s Yq12 length relative to the global or haplogroup-specific median. Fisher’s exact tests identified significant associations for both comparisons, with a stronger association when paternal Yq12 length was evaluated relative to the haplogroup-specific median than the global median (*haplogroup-specific*: Fisher’s exact p-value = 1.402e⁻^5^; *global*: Fisher’s exact p-value = 0.012, **Supplementary Table S13**). Fathers carried the longer estimated Yq12 in 81.2% of pairs (134/165) when paternal length was above the haplogroup-specific median, compared with 58.0% (80/138) when it was below; the corresponding values for the global median were 77.6% (111/143) and 64.4% (103/160). Sons were therefore more likely to carry the longer Yq12 when their fathers had relatively short Yq12 regions (odds ratio for father longer, below versus above median: 0.320, 95% CI 0.184-0.551 for the *haplogroup-specific* median; 0.522, 95% CI 0.302-0.892 for the *global* median), though fathers carried the longer Yq12 in most pairs in both groups (**Figure 3c**). We also tested whether the directional asymmetry observed for Yq12 length extended to repeat composition, given the clustering of *DYZ1*:*DYZ2* ratios near parity across males. Across 303 father-son pairs, the son’s log-transformed *DYZ1:DYZ2* ratio was closer to zero (1:1 ratio) than the father’s in 61.4% of pairs (186/303; 95% CI: 55.6–66.9%; one-sided sign test, p-value=4.39e^−5^; **Figure 3d**), supporting a tendency for sons to be closer to parity than their fathers. Across both Yq12 length and repeat composition, father-son differences showed a directional asymmetry, with sons tending toward shorter arrays and *DYZ1*:*DYZ2* ratios closer to parity.

### Yq12 Expression

The remarkable size variability of Yq12, together with the generally near-balanced *DYZ1*:*DYZ2* copy-number ratios and previous reports of *DYZ1*-derived transcripts in testis^34^, motivated us to investigate Yq12 expression in human testis. Using LRKmer to screen a pooled testis-derived PacBio RNA Iso-Seq long-read dataset (PRJNA659539)^56^, we identified reads containing partial or full-length *DYZ1* and *DYZ2* repeat units, including reads spanning more than one repeat unit (**Supplementary Table S14)**. These data provide evidence for both *DYZ1*- and *DYZ2*-derived transcription in human testis and production of multi-repeat Yq12-derived RNA molecules.

To further investigate the origin of Yq12 transcripts, we next screened a human testis scRNA-seq dataset of three male donors across testicular cell types from the adult human testis transcriptional cell atlas^57^. Using the k-mer framework underlying SRKmer, we found that reads containing *DYZ1*- or *DYZ2*-specific k-mers were most abundant in male germ cells, particularly haploid spermatids (**Figure 4a; Supplementary Table S15**). A more refined stage-specific analysis across spermatogenesis revealed marked changes in Yq12-derived read abundance during germ-cell differentiation, with round spermatid stages 4 and 5 showing the highest levels of both *DYZ1* and *DYZ2* (**Figure 4b,c; Supplementary Table S16**).

**Figure 4:**
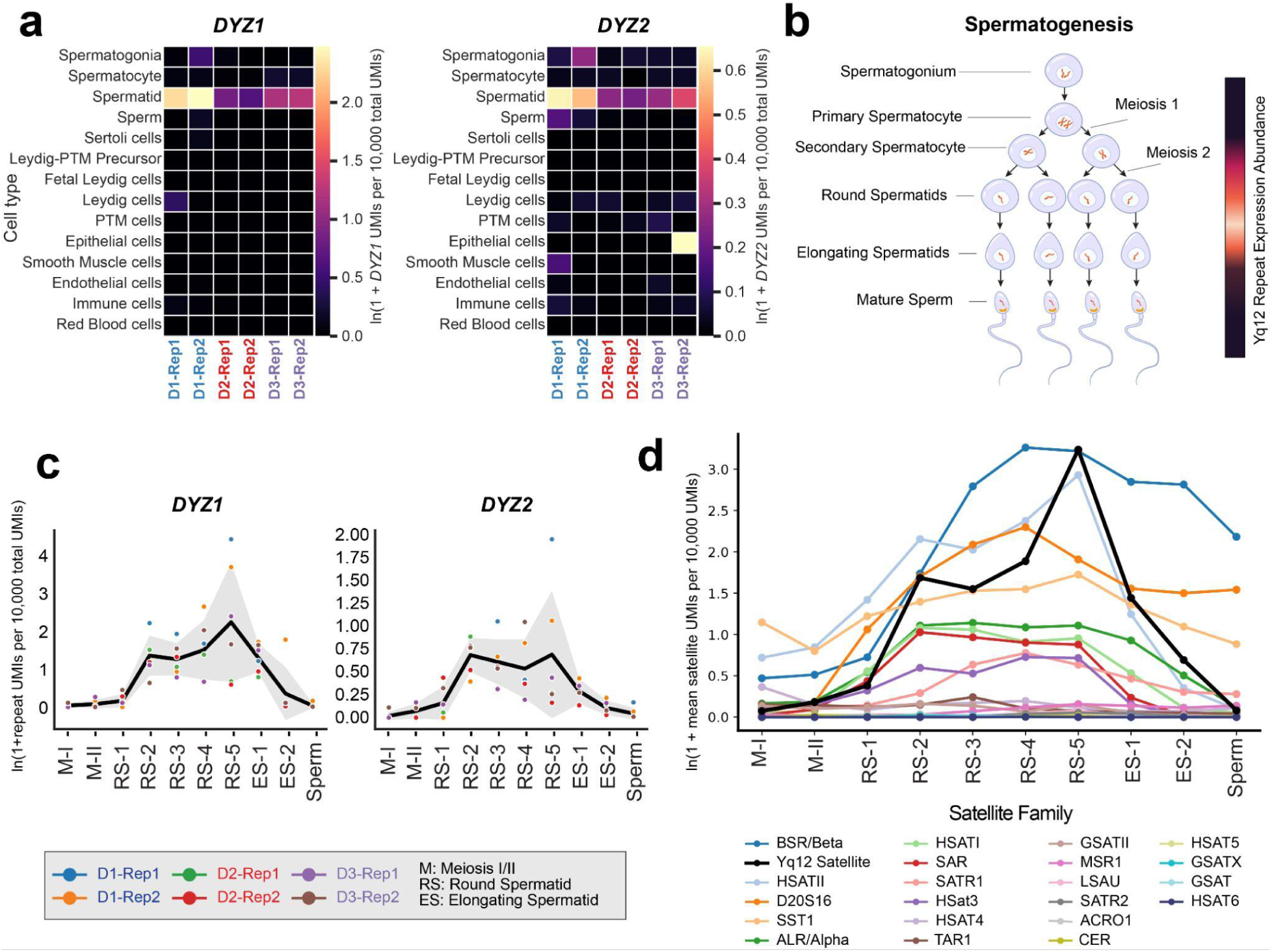
Yq12 satellite-containing transcripts during human spermatogenesis. **a)** Heatmap of natural log-transformed *DYZ1*- and *DYZ2*-specific UMI abundance across somatic and germ cell types in adult testes from three donors (D1: 17 years old; D2: 24 years old; D3: 25 years old), each with two replicates. Values represent ln(1 + *DYZ1* or *DYZ2* UMIs per 10,000 total UMIs). **b)** Schematic of spermatogenesis, with relative Yq12 satellite expression indicated from low (dark purple) to high (white). **c)** Expression dynamics of *DYZ1*- and *DYZ2*-specific UMIs from meiosis I through mature sperm across the three donors. Values represent ln(1 + repeat-specific UMIs per 10,000 total UMIs). Colored points represent individual donor replicates at each germ-cell stage, and the black line represents the mean of these values at each stage. **d)** Normalized abundance of transcripts containing different satellite sequences across male germ-cell development. For each satellite family, UMI abundance was calculated across the indicated germ-cell stages and plotted as ln(1 + mean satellite UMIs per 10,000 total UMIs), with means calculated across donor replicates. *DYZ1* and *DYZ2* were combined as Yq12 satellite, which showed a stage-dependent pattern with the highest abundance in late round spermatids, particularly RS-5.

To assess whether Yq12 transcription reflected a broader increase in satellite-derived RNA during spermatogenesis, we additionally quantified transcripts containing other satellite families using RepeatMasker (**Figure 4d; Methods**). Satellite families showed heterogeneous developmental profiles, with peak abundance occurring at different stages of germ-cell differentiation rather than a uniform increase across families. Among non-Yq12 satellites, HSATII showed a strong and reproducible correspondence with Yq12-derived transcripts across development. Donor-mean developmental profiles correlated strongly (Pearson’s r = 0.905; **Supplementary Table S17**), and stage-to-stage changes in HSATII abundance correlated positively with changes in Yq12-derived satellite reads within each donor (r = 0.985, 0.490, and 0.853 for D1–D3, respectively; **Supplementary Table S17**). HSATII and Yq12-derived satellite abundance also changed synchronously across 7 of 9 developmental transitions in all three donors (**Supplementary Table S17**). Similarly, Z-score-normalized trajectories showed coordinated increases during round-spermatid differentiation, followed by declines during elongating spermatid stages and sperm (**Supplementary Figure S6**). These results show that Yq12-derived transcripts exhibit developmentally structured expression during human spermatogenesis, reaching their highest abundance in haploid round spermatids (RS-5), while broader satellite transcription follows family-specific rather than uniformly elevated developmental patterns.

## Discussion

Yq12 remains one of the most technically challenging parts of the human genome to assemble, measure, and compare between males^5–7^. This study shows that assembly-informed approaches can reliably estimate Yq12 length and repeat composition from sequencing reads, avoiding the need to generate a complete Y-chromosome assembly for each individual. We first benchmarked two complementary approaches, SRKmer and LRKmer, against complete Yq12 assemblies, with both showing strong agreement for Yq12 length estimation. We then used SRKmer for the population-scale analyses of Yq12 length and *DYZ1*:*DYZ2* composition, while LRKmer provided an independent long-read estimator and enabled Yq12 sequence detection in long-read transcription datasets. These results extend previous alignment-free approaches that focused primarily on *DYZ1*^8^ and demonstrate that the full Yq12 region can be treated as a measurable quantitative genomic trait. Our assembly-independent framework therefore enables population-scale investigation of Yq12 variation.

Across 1,597 males from the 1kGP, Yq12 emerged as a highly variable (less than 1 Mb to greater than 50 Mb) but evolutionarily structured component of the Y chromosome. Size differences among major Y-chromosome haplogroups are consistent with repeat expansions and contractions accumulating in non-recombining paternal lineages, allowing lineage-specific Yq12 architectures to develop over time^6,15,23^. At the same time, the wide variation observed within haplogroups shows that Y-chromosome ancestry is not the only factor shaping Yq12 length. Beyond this lineage-specific structure, phylogenetic model comparisons indicate that Yq12 length does not simply accumulate through unconstrained evolutionary drift. Instead, an Ornstein–Uhlenbeck process describes Yq12 length substantially better than Brownian motion, with an inferred long-term equilibrium near 26 Mb and a broad inferred stationary range of approximately 16-44 Mb. These results are consistent with Yq12 size evolving within a broad but constrained range. Such mean-reverting dynamics could arise through stabilizing selection, mutational biases, or repeat-associated mechanisms that shape the balance between array expansion and contraction; the evolutionary models cannot distinguish among these possibilities^54,70,71^.

We observed a parallel pattern of long-term constraint in Yq12 repeat composition. Despite a >50-fold range in estimated Yq12 length, *DYZ1* and *DYZ2* abundance remained closely balanced across males. Our findings further support a near-balanced relationship between *DYZ1* and *DYZ2*^5,6^. Furthermore, our phylogenetic analyses provide evidence that this pattern reflects evolutionary constraint on repeat composition. *DYZ1* and *DYZ2* showed strongly correlated evolutionary change, with substantially greater variation in their shared abundance than in their relative imbalance. Moreover, a stationary OU process explained *DYZ1*:*DYZ2* composition better than unconstrained Brownian evolution, while allowing the inferred equilibrium to differ from parity did not significantly improve model fit. Together, these findings support long-term evolutionary constraint around a shared near-parity equilibrium despite extensive variation in overall Yq12 size. However, the balance was not completely invariant. Larger Yq12 regions contained proportionally more *DYZ2*, consistent with the greater evolutionary variance observed for *DYZ2* than *DYZ1*. This constraint coexists with strong phylogenetic structure, indicating that lineage-specific deviations in *DYZ1*:*DYZ2* composition can persist without requiring distinct haplogroup-specific equilibria. Importantly, the branchwise analysis did not detect an excess of evolutionary changes directed toward parity, suggesting that the phylogenetic data do not provide evidence for systematic branchwise restoration toward a 1:1 ratio. Instead, the combination of coordinated *DYZ1* and *DYZ2* evolution and long-term constraint around near parity indicates that balanced repeat composition can be maintained despite individual evolutionary changes that are not consistently directed toward parity.

The father-son analyses provide a complementary view of Yq12 variation over much shorter timescales. Across more than 300 father-son pairs, sons tended to have shorter estimated Yq12 regions than their fathers and more often had *DYZ1:DYZ2* ratios closer to 1:1. Although these observations are directionally consistent with the long-term constraints on Yq12 size and composition inferred from the phylogenetic analyses, they do not establish a directed process that restores Yq12 toward a particular size or repeat ratio during transmission. Several mechanisms could contribute to the observed asymmetry. Repeat restructuring within the paternal germline could generate Yq12 variants that differ from those measured in paternal somatic DNA, while age-associated somatic changes could cause the sampled paternal Yq12 to differ from the array present in the germline. We also cannot exclude selective transmission or survival of particular Yq12 configurations; for example, if germline variation in Yq12 architecture exists, differences in germ-cell or sperm success, fertilization, or subsequent developmental viability could bias which configurations are observed in sons. Such germline variation would be consistent with the high mutation rate previously measured in Yq12^6,23^ and with the propensity of large tandem-repeat arrays to expand or contract through unequal exchange and related recombination-based mechanisms. With the available lymphoblastoid-cell-line data, however, we cannot distinguish among germline restructuring, somatic variation, and selective transmission or survival. The asymmetry also varied across pairs. Sons more often carried the longer Yq12 when their fathers fell below the median, particularly the father’s haplogroup-specific median, than when fathers were above it. This pattern could reflect lineage-specific Yq12 architecture^6^ and repeat-mediated turnover, but it could also be influenced by regression toward the mean or estimation error^72^.

The long-term constraint observed for both Yq12 size and *DYZ1*:*DYZ2* composition raises the possibility that Yq12 has biological relevance beyond being a passive heterochromatic block, although its functional significance remains poorly understood. Yq12 has traditionally been regarded as constitutive heterochromatin^16^, and substantial variation in its size is compatible with an apparently normal phenotype^5,6^. However, extreme Yq12 configurations and Yq12-associated chromosomal rearrangements have been reported in association with male infertility, azoospermia, meiotic arrest, and recurrent pregnancy loss, suggesting potential biological consequences in at least some contexts^18,26,27^. Our findings add two observations that further strengthen that Yq12 has biological relevance. First, Yq12 size and *DYZ1*:*DYZ2* composition showed nonrandom population and evolutionary structure, including evidence for long-term constraint on both total Yq12 length and repeat composition. Second, we detected reads containing *DYZ1*- and *DYZ2*-derived k-mers in testis transcriptomic data. Using our long-read k-mer estimator, LRKmer, on testis Iso-Seq data^56^, we identified long reads containing partial or full *DYZ1* and *DYZ2* repeat units, including reads spanning more than one repeat unit. To our knowledge, *DYZ2*-derived testis transcripts have not previously been described. We then applied the short-read k-mer framework, SRKmer, to human testis scRNA-seq data^57^ and discovered that reads containing *DYZ1*- and *DYZ2*-specific k-mers were enriched in male germ cells, particularly haploid spermatids, with the highest abundance in round spermatid stages 4 and 5.

Comparison with other satellite families showed that satellite transcription was not uniformly elevated during spermatogenesis; instead, individual satellite families followed different developmental trajectories. Notably, HSATII showed a particularly strong and reproducible relationship with Yq12, including correlated stage-to-stage changes during spermatogenesis across all three donors. This similarity indicates that the Yq12 pattern is not unique, but the heterogeneous behavior of other satellite families argues against a nonspecific global increase in satellite transcription. Rather, these findings are consistent with selective, developmentally structured transcription of satellite DNA during male germ-cell differentiation. The coordinated developmental profiles of Yq12- and HSATII-derived transcripts are particularly notable because HSATII is a pericentromeric satellite known to undergo regulated transcription in other contexts^73,74^. DUX4, a transcription factor normally expressed in human testis^75^, directly binds HSATII and can activate its transcription^76^. Prior work has also documented transcriptional activity of pericentromeric heterochromatin during specific stages of meiosis^77^. These observations provide precedent for regulated satellite transcription in the male germ line and raise the possibility that distinct satellite families respond to shared developmental or transcriptional programs during spermatid differentiation. However, the present data do not establish a common regulatory mechanism or functional role for these transcripts.

The enrichment of Yq12-derived transcripts in round spermatids raises the possibility that Yq12 transcription is linked to chromatin remodeling during spermiogenesis. In *Drosophila*^78^, transcription through AAGAG satellite DNA in spermatocytes facilitates heterochromatin remodeling in preparation for the histone-to-protamine transition, even though the resulting satellite RNA itself appeared dispensable. Notably, the severity of subsequent sperm chromatin-compaction defects depended on the amount of AAGAG satellite DNA present. Future studies will need to determine whether transcription through the large and highly variable human Yq12 satellite arrays contributes to remodeling of Yq12 heterochromatin during sperm chromatin packaging. Given the extensive inter-individual variation in Yq12 size identified here, this model raises the possibility that Y-bearing spermatids from different males vary substantially in the amount of Yq12 heterochromatin that must undergo remodeling during spermiogenesis, a period of extensive chromatin reorganization^57^ that coincides with peak Yq12-derived transcript abundance.

Beyond a potential role in chromatin remodeling, Yq12 variation could also have regulatory consequences through a distinct, sequence-dependent mechanism. HSat3 arrays have been proposed to act as megabase-scale transcription-factor binding platforms, carrying millions of motif copies bound by more than a dozen factors from diverse signaling pathways^36^. *DYZ1* corresponds to the Y-specific HSat3A6 subfamily^4,8^, which is sequence-divergent and exhibits a distinct motif-enrichment profile, including enrichment for BCL11B, CDX4, ETV5::FOXO1, and HOXD9 motifs^36^. If satellite arrays function as reservoirs of transcription-factor binding sites, then variation in *DYZ1* array size could translate into substantial differences in the number of available binding sites among males. Any such effect would likely depend on *DYZ1* content rather than total Yq12 length alone, because Yq12 also contains interspersed *DYZ2*/HSat1 blocks^4^, and HSat1 lacks the motif density that characterizes HSat3 for at least some of these factors^36^. However, this possibility remains speculative: no HSat3-dependent regulatory mechanism has been demonstrated specifically at Yq12, and array length has not been shown to scale with transcription-factor recruitment. Locus-specific localization and perturbation experiments will therefore be needed to determine whether transcription-factor recruitment links Yq12 variation to germ-cell biology.

There are several limitations to this study. First, although the SRKmer performed strongly in benchmarking, it remains an indirect estimator of Yq12 length. Extreme estimates, particularly very short Yq12 calls, require careful interpretation and ideally warrant validation using orthogonal long-read assembly or cytogenetic approaches. Second, LRKmer benchmarking showed good agreement with assembled Yq12 lengths, but performance was less consistent than that of SRKmer, potentially reflecting systematic HiFi coverage biases previously observed across specific human satellite arrays^2,79^. Third, we derived Yq12 estimates for both fathers and sons from lymphoblastoid cell lines, which may not represent the germline arrays transmitted between generations. Data on paternal age and multiple tissues, which would allow somatic and germline contributions to be separated, were not available. In addition, the observed relationship between paternal Yq12 length and father-son differences may be influenced by measurement error or regression toward the mean^72^. The observed directional patterns should therefore be confirmed in larger pedigrees with complete Yq12 assemblies. Finally, although the expression analyses detected reads containing Yq12-derived k-mers, the short-read 10x data have limited ability to resolve transcripts from highly repetitive satellite arrays. Consequently, we cannot always assign these reads to precise positions within Yq12, nor can they establish whether *DYZ1*- and *DYZ2*-containing reads originate from full-length, stable, or functional transcripts.

Despite these limitations, the ability to estimate Yq12 length from standard short-read sequencing opens several important applications. For genome assembly, sample-specific Yq12 estimates can provide an expected target size for one of the most frequently collapsed or underrepresented regions of the Y chromosome^5,6^, helping guide assembly evaluation and reducing uncertainty when Yq12 cannot be fully resolved. For population genetics, Yq12 length becomes a measurable Y-linked structural trait that can be analyzed across haplogroups, populations, and pedigrees. Biomedical studies can now incorporate Yq12 length and *DYZ1*:*DYZ2* composition as covariates in analyses of male infertility, recurrent pregnancy loss, age-related mosaic loss of the Y chromosome, immune phenotypes, cancer risk, and other traits to which Y-chromosome variation may contribute^24–27,80,81^. Population-scale analyses across more diverse cohorts will further clarify how demographic history and Y-lineage architecture shape Yq12 size distributions. Future studies should determine whether variation in Yq12 size, repeat composition, transcription, or chromatin state has functional consequences for germ-cell development or other Y-linked phenotypes.

In summary, SRKmer and LRKmer offer a scalable way to measure one of the largest and hardest-to-access heterochromatic regions in the human genome. The population data revealed wide variation in Yq12 length that is structured by Y-chromosome lineage yet constrained over longer evolutionary timescales, while *DYZ1* and *DYZ2* showed coordinated evolution around a shared near-parity equilibrium. The father-son and transcriptomic analyses reveal testable patterns of intergenerational change and germ-cell transcription, while identifying the validation needed to establish the underlying mechanisms. Overall, these findings establish Yq12 as a dynamic, compositionally constrained, and transcriptionally active component of the human Y chromosome and provide the tools needed to investigate its evolutionary and biological consequences.

## Acknowledgments

We thank the individuals who contributed samples to the 1kGP, the Human Pangenome Reference Consortium (BioProject PRJNA730823) and its funder, the National Human Genome Research Institute, and the Human Genome Structural Variation Consortium. We also thank the staff at Clemson University for providing computational resources through the Palmetto HPC and The Jackson Laboratory for Genomic Medicine for providing computational resources through the Sumner HPC. During preparation of this work, the authors used ChatGPT (OpenAI) and Claude (Anthropic) to assist with aspects of code development and troubleshooting, grammar verification, and language refinement of selected portions of the manuscript. All AI-assisted content was reviewed, edited, and verified by the authors, who take full responsibility for the final manuscript and analyses.

## Author Contributions

Mark Loftus (M.L.): Conceptualization, Data generation, Formal analysis, Methodology, Validation, and Writing. Pille Hallast (P.H.): Formal analysis, Writing, and Visualization. Gianni V. Martino (G.V.M.): Data generation. Gianluca Petris (G.P.): Writing and Formal analysis. Charles Lee (C.L.): Writing and Formal analysis. Miriam K. Konkel (M.K.K.): Conceptualization, Methodology, Writing, and Formal analysis.

## Supplementary Data Statement

Supplementary Data are available online.

## Code and Data Availability

The data underlying this article are publicly available, with accession numbers and source information provided in the article and its online supplementary material. Source code for the Yq12 estimation tools developed in this study is available through Zenodo at https://doi.org/10.5281/zenodo.23219520 and on GitHub at https://github.com/Markloftus/Yq12Estimations.

