## Supplemental Figures for "Assembly-independent measurement of human Yq12 heterochromatin reveals population-scale variation and Yq12-derived RNA in testis"

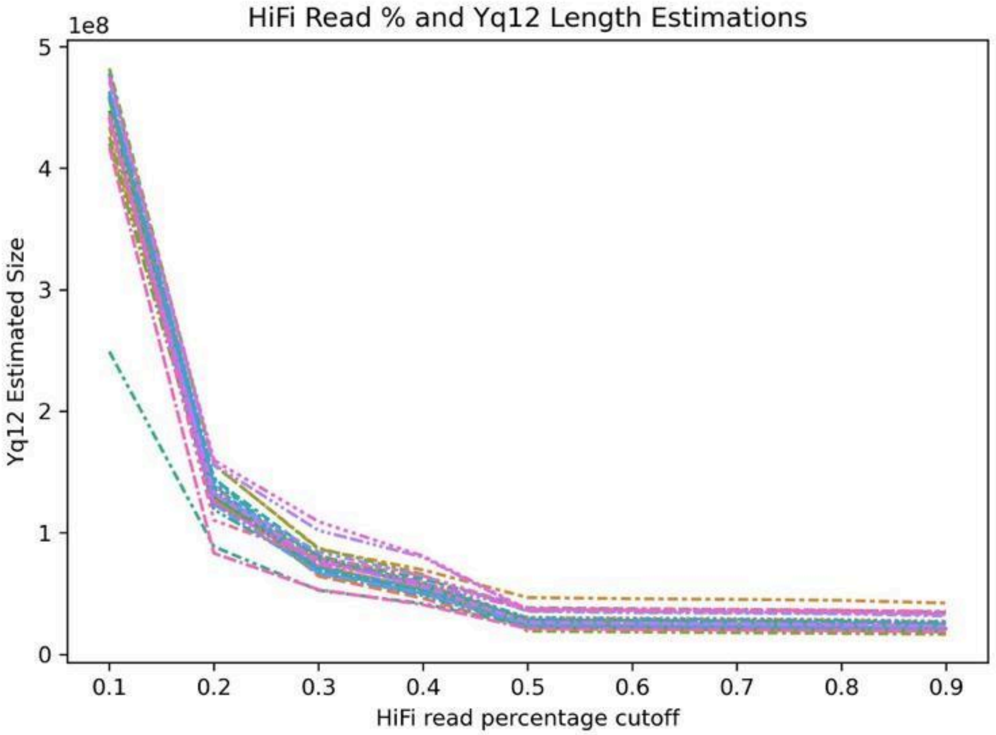


**Supplemental Figure 1) HiFi sample Yq12 length estimations at different read percentage cutoffs**
A lineplot showing the estimated Yq12 length (y-axis) at different read percentage cutoffs (x-axis). Across samples the LRKmer derived Yq12 estimated length appears to level off at the 50% of read percentage cutoff.

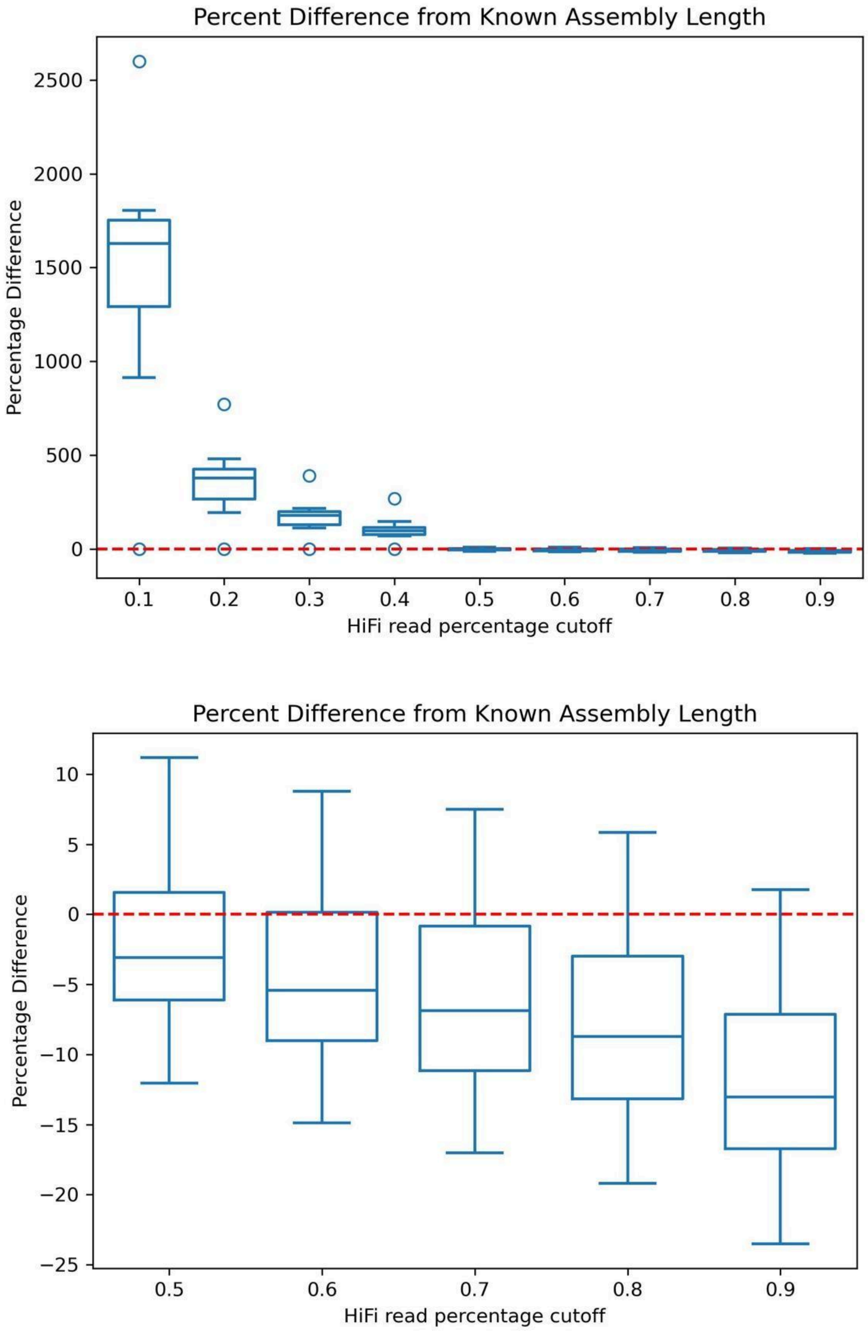


**Supplemental Figure 2) LRKMer read kmer profile cutoffs**
**Top panel:** Boxplots showing the percent difference between LRKmer-estimated and *de novo* assembled Yq12 lengths across 74 samples at different read-profile percentage cutoffs. Agreement with the assembled lengths was greatest at the 50% cutoff. **Bottom panel:** Magnified view of the same data across the 50%–90% HiFi read-percentage cutoffs. At cutoffs above 50%, LRKmer increasingly underestimated Yq12 length, with a mean underestimation exceeding 5%.

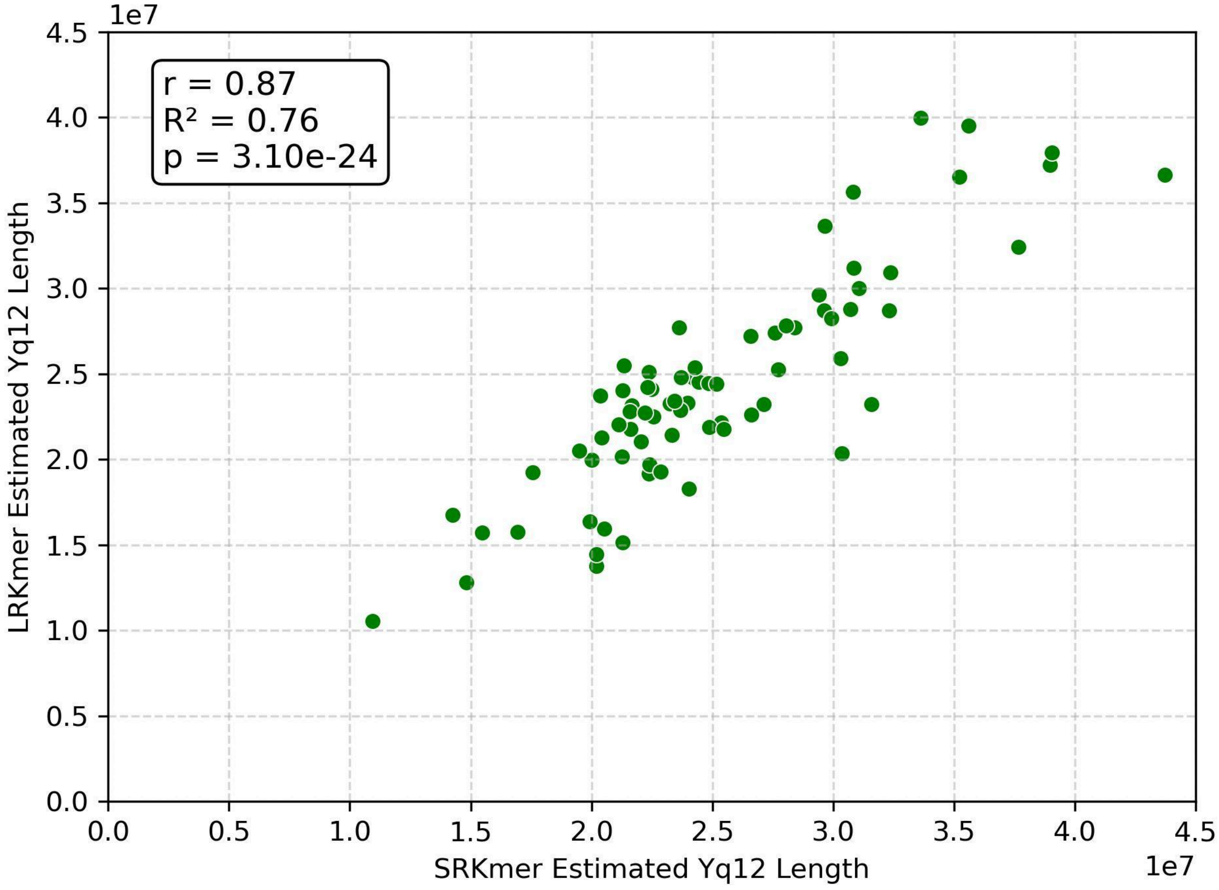


**Supplemental Figure 3) LRKmer estimated vs SRKmer estimated Yq12 lengths**

A scatterplot showing a strong correlation (Pearson r=0.874) between the SRKmer estimated Yq12 length (x-axis) and the LRKmer estimated Yq12 length (y-axis) in 74 male samples.

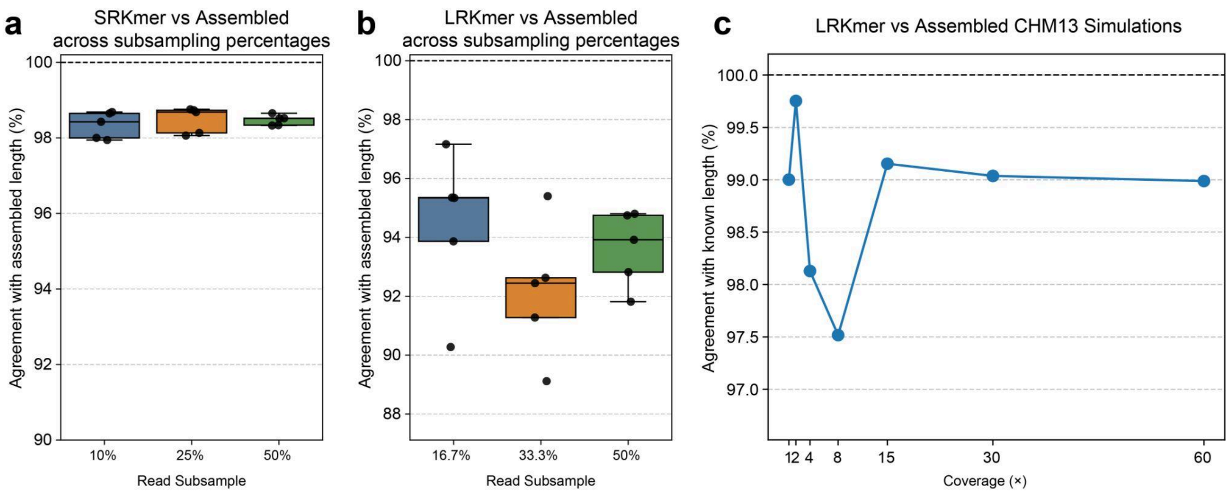


**Supplemental Figure 4) SRKmer and LRKmer Subsampling Benchmarking**

**a)** Agreement (accuracy see **methods**) between SRKmer-estimated and assembled Yq12 lengths following subsampling of the HG01890 short-read dataset, which had approximately 15.5x Y-chromosome coverage before subsampling. Reads were independently subsampled five times at 10%, 25%, and 50% of the original dataset. Points represent individual subsampling replicates, and boxplots summarize the distribution at each subsampling level. SRKmer estimates remained highly consistent across subsampling depths and recovered approximately 98–99% of the assembled Yq12 length, indicating a small but consistent underestimation. **b)** Agreement between LRKmer-estimated and assembled Yq12 lengths following subsampling of the randomly selected male, HG01890, long-read dataset, which had approximately 6x Y-chromosome coverage before subsampling. Reads were independently subsampled five times at 16.7%, 33.3%, and 50% of the original dataset. LRKmer estimates recovered approximately 89–97% of the assembled Yq12 length, with the greatest underestimation observed among some lower-coverage subsamples. **c)** Agreement between LRKmer estimates and the known CHM13 Yq12 length using simulated PacBio HiFi whole-genome read datasets generated at 1x, 2x, 4x, 8x, 15x, 30x, and 60x coverage. LRKmer recovered approximately 97.5–99.8% of the known Yq12 length across all simulated coverage levels. The dashed horizontal line at 100% represents perfect agreement with the assembled or known Yq12 length. Together, these analyses show that SRKmer and LRKmer estimates are generally robust to variation in read depth, although LRKmer displayed greater variability at lower coverage.

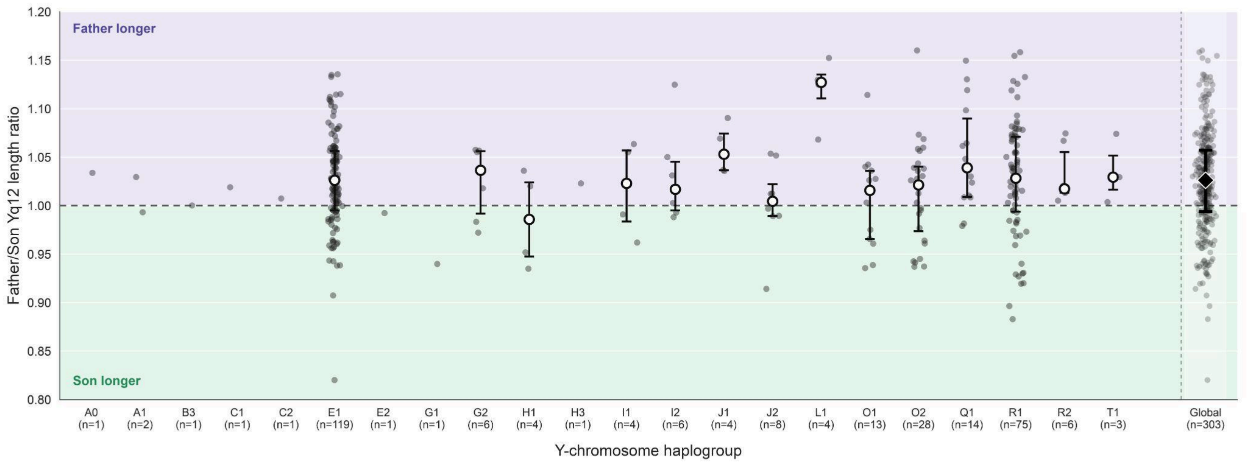


**Supplemental Figure 5) Father-to-son Yq12 length ratios across Y-chromosome haplogroups.** Each gray point represents an individual father-son pair, plotted as the ratio of the father's estimated Yq12 length to that of his son. Ratios greater than 1 indicate a longer Yq12 in the father, whereas ratios less than 1 indicate a longer Yq12 in the son; the horizontal dashed line marks equal father and son lengths (ratio = 1). Open circles indicate the median ratio within each haplogroup, with vertical bars spanning the interquartile range (25th-75th percentiles) for haplogroups containing at least three pairs. Sample sizes are shown below each haplogroup. The black diamond denotes the median across all 303 father-son pairs included in the primary analysis, with the vertical bar indicating the interquartile range; the global summary is separated from the haplogroup-specific estimates by the vertical dotted line. Six pairs with father-son Yq12 length differences ≥20% were excluded from the primary analysis. Background shading indicates regions in which the father (purple) or son (green) has the longer estimated Yq12.

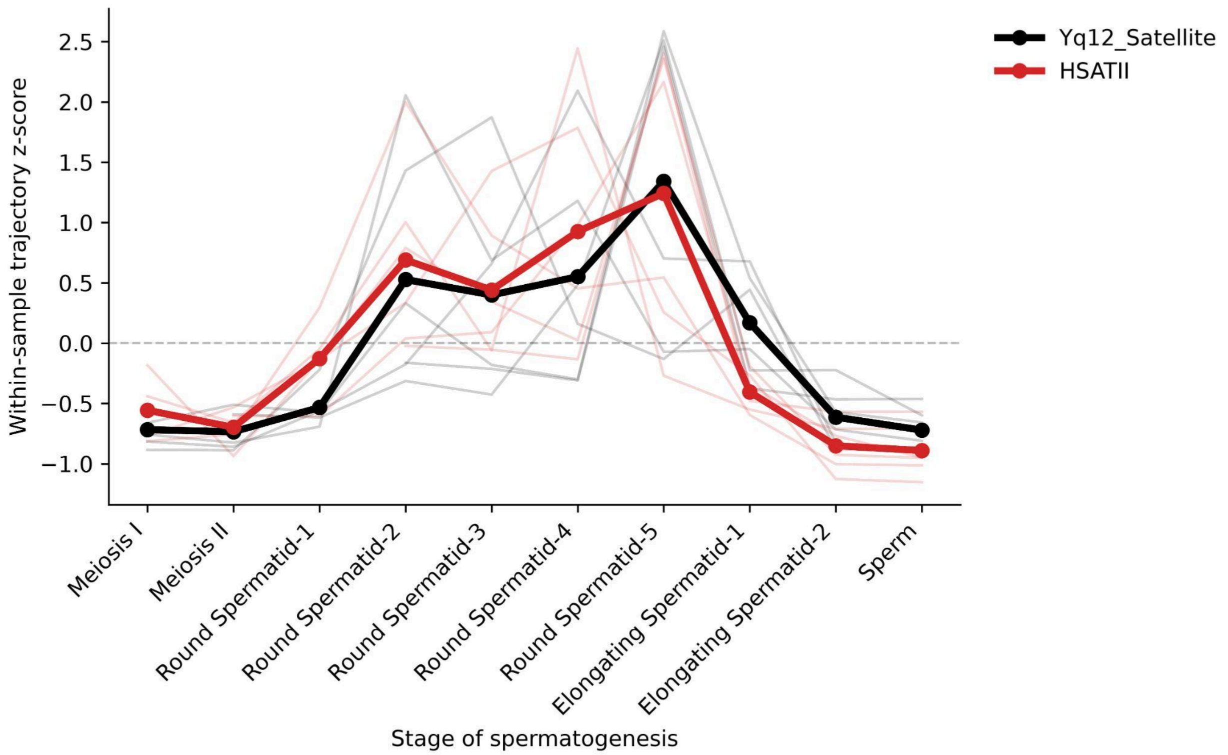


**Supplemental Figure 6) Developmental trajectories of Yq12 and HSATII transcription during spermatogenesis.** Yq12 (*DYZ1* + *DYZ2*) and HSATII UMI abundances were z-score normalized within each sample to compare developmental trajectory shape independently of absolute abundance. Light lines show individual sample trajectories, and thick lines show the mean trajectory across samples. Both satellite families increase during round-spermatid differentiation, reach their highest mean values in late round spermatids, and decline during elongating spermatid stages and sperm.
